# Dissecting the respective roles of microbiota and host genetics in the susceptibility of *Card9*^-/-^ mice to colitis

**DOI:** 10.1101/2023.01.03.522555

**Authors:** C. Danne, B. Lamas, A. Lavelle, M-L. Michel, G. Da Costa, Hang-Phuong Pham, A. Lefevre, C. Bridonneau, M. Bredon, J. Planchais, M. Straube, P. Emond, P. Langella, H. Sokol

**Author notes:** Authors share co-first authorship (C. Danne, B. Lamas). Corresponding Author: Camille Danne, PhD, CRSA Hôpital Saint-Antoine, 27 rue de Chaligny, 75012 Paris, France; Harry Sokol, MD, PhD, Service de Gastro-entérologie, Hôpital Saint-Antoine, 184 rue du Faubourg Saint-Antoine, 75571 Paris Cedex 12, France.

## Abstract

**Background:** The etiology of Inflammatory Bowel Disease (IBD) is unclear but involves both genetics and environmental factors, including the gut microbiota. Indeed, exacerbated activation of the gastrointestinal immune system toward the gut microbiota occurs in genetically susceptible hosts and under the influence of the environment. For instance, a majority of IBD susceptibility loci lie within genes involved in immune responses, such as caspase recruitment domain member 9 (*Card9*). However, the relative impacts of genotype versus microbiota on colitis susceptibility in the context of CARD9 deficiency remain unknown.

**Results:** *Card9* gene directly contributes to recovery from dextran sodium sulfate (DSS)-induced colitis by inducing the colonic expression of the cytokine IL-22 and the antimicrobial peptides *Reg3β* and *Reg3γ* independently of the microbiota. On the other hand, *Card9* is required for regulating the microbiota capacity to produce AhR ligands, which leads to the production of IL-22 in the colon, promoting recovery after colitis. In addition, cross-fostering experiments showed that five weeks after weaning, the microbiota transmitted from the nursing mother before weaning had a stronger impact on the tryptophan metabolism of the pups than the pups’ own genotype.

**Conclusions:** These results show the role of CARD9 and its effector IL-22 in mediating recovery from DSS-induced colitis in both microbiota-independent and microbiota-dependent manners. *Card9* genotype modulates the microbiota metabolic capacity to produce AhR ligands, but this effect can be overridden by the implantation of a WT or “healthy” microbiota before weaning. It highlights the importance of the weaning reaction occurring between the immune system and microbiota for host metabolism and immune functions throughout life. A better understanding of the impact of genetics on microbiota metabolism is key to developing efficient therapeutic strategies for patients suffering from complex inflammatory disorders.

## Introduction

Inflammatory bowel diseases (IBD), including Crohn’s disease and ulcerative colitis, are characterized by chronic pathological inflammation of the digestive tract. The pathogenesis of IBD remains unclear but involves activation of the gastrointestinal immune system toward the gut microbiota in genetically susceptible hosts and under the influence of the environment ^1^. The gut microbiota, composed of bacteria, fungi, and other microorganisms is fundamental to the health and nutrition of the host ^2^. Loss of the fragile equilibrium within this complex ecosystem (termed dysbiosis), which is often characterized by a decreased biodiversity, overgrowth of potentially harmful microorganisms, and disappearance of protective ones, can trigger numerous pathologies, including IBD ^3^. Genetic factors are also associated with IBD pathogenesis and a majority of IBD susceptibility loci lie within genes involved in immune responses, such as caspase recruitment domain member 9 (*Card9*).

*Card9* is highly expressed in myeloid cells, including dendritic cells, macrophages and neutrophils, and encodes an adaptor protein that integrates signals downstream of pattern recognition receptors. CARD9 is involved in the immune response to fungi, mycobacteria and bacteria ^4,5^. It acts on several pathways, such as NF-kB and p38/Janus Natural Kinase, and modulates Toll-like receptor signaling, inducing cytokine production and immune cells activation, leading to the elimination of detected microorganisms ^4–6,7^. In a previous study, we showed that CARD9 mediates recovery from colitis through the production of IL-22, a cytokine with well-known effects on intestinal homeostasis and barrier function ^8–10^. *Card9*^-/-^ mice have an enhanced susceptibility to dextran sodium sulfate (DSS)-induced colitis and an increased load of gut-resident fungi. Moreover, we also demonstrated that the transfer of *Card9*^-/-^ mice microbiota to wild-type (WT) germ-free (GF) recipients was sufficient to recapitulate the defective IL-22 activation and the increased colitis susceptibility observed in *Card9*^-/-^ mice ^8^. This defect was due to the impaired ability of bacterial microbiota from *Card9*^-/-^ mice to metabolize tryptophan (Trp) into aryl hydrocarbon receptor (AhR) ligands, such as indole derivatives ^8^. Indeed, recent data have indicated that Trp catabolites from the microbiota play a role in mucosal immune responses via AhR, which in turn modulates the production of IL-22 ^11,12^. In humans, comparable mechanisms appear to be involved, as we showed that the microbiota of patients with IBD exhibits impaired production of AhR ligands, which correlates with IBD-associated single-nucleotide polymorphisms (SNP) within CARD9 ^8^. However, the circular causality concept in which alterations in the microbiota fuel intestinal inflammation that, in turn, worsens microbiota alterations remains to be deeply explored.

Here we demonstrated that *Card9* modulates the susceptibility to DSS-induced colitis in both microbiota-dependent and microbiota-independent manners. Using GF *Card9*^-/-^ mice, we showed that *Card9* promotes the colonic expression of the cytokine IL-22 and the antimicrobial peptides REG3β and REG3γ independently of the microbiota. On the other hand, colonization of GF *Card9*^-/-^ mice with a WT microbiota showed that the *Card9* gene is required for shaping the bacterial microbiota composition and regulating its capacity to produce AhR ligands. In addition, cross-fostering experiments showed that the microbiota transmitted from the nursing mother has a stronger impact on Trp metabolism and AhR activity at adult age, than has the pups’ own genotype. These results show the role of CARD9 and its effector IL-22 in mediating recovery from colitis in both microbiota-independent and microbiota-dependent manners, and highlight the importance of the weaning reaction occurring between the immune system and microbiota for host metabolism and immune functions throughout life.

## Results

### *Card9* contributes to colitis recovery and controls intestinal immune response independently of the gut microbiota

We previously showed that *Card9*^-/-^ mice microbiota contributes to susceptibility to DSS-induced colitis^8^. However, the specific contributions of the genetic, i.e. the deletion of *Card9*, independently of the microbiota, remains to be explored. We first questioned whether *Card9* gene does contribute to colitis recovery in a gut microbiota-independent manner by exposing GF WT and GF *Card9*^-/-^ mice to DSS. Compared to GF WT mice, recovery was impaired in GF *Card9*^-/-^ mice after DSS-induced colitis, with delayed weight gain, higher disease activity index, and greater histopathologic alterations (Fig. 1A-C). To examine the mechanisms responsible for this defect, we compared the colon transcriptomes of GF WT and GF *Card9*^-/-^ during the course of the colitis. Compared to GF WT mice, GF *Card9*^-/-^ exhibited a significant downregulation of genes involved in host defense, as well as in immune and inflammatory responses at day 7 (Fig 1D), supporting a defective intestinal immune response. These results were confirmed by real-time quantitative PCR in colon tissue showing that *Il-22, Reg3β* and *Reg3γ* expression was induced at day 7 and 12 in GF WT mice but not in GF *Card9*^-/-^ mice (Fig. 1E). Moreover, the expression of *Il1β* and *Tnfα* was decreased in the colon of GF *Card9*^-/-^ mice compared to GF WT mice at day 7 (Fig. 1F). At day 12, *Tnfα* expression was significantly higher in GF *Card9*^-/-^ mice compared to GF WT mice, which could reflect the delayed colitis recovery in the absence of *Card9* (Fig. 1F). This defect in *Il-22* expression, a cytokine known to regulate mucosal wound healing^13^, associated with the increased expression of the pro-inflammatory cytokines *Il1β* and *Tnfα*, may play a role in the higher colitis susceptibility of GF *Card9*^-/-^ mice. These results demonstrate that *Card9* contributes to colitis recovery in a gut microbiota-independent manner by supporting the normal intestinal immune response.

**Figure 1.**
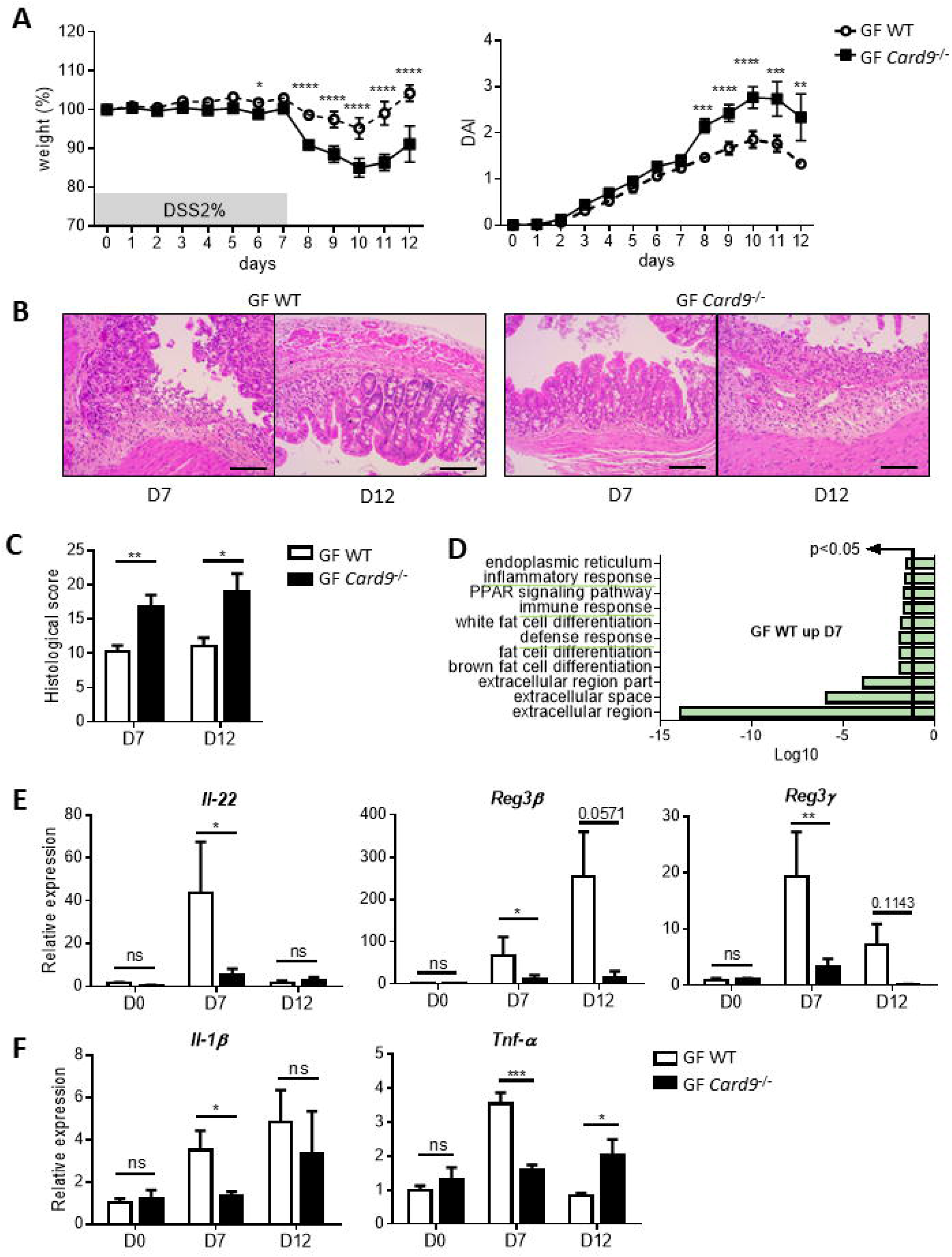
*Card9* contributes to colitis recovery and controls intestinal immune response independently of the gut microbiota. (A) Weight (left) and Disease Activity Index (DAI, right) score of DSS-exposed GF WT and GF *Card9*^-/-^ mice. (B) Representative H&E-stained images of colon cross-sections from DSS-exposed GF WT (upper panel) and GF *Card9*^-/-^ (lower panel) mice at day 7 (left) and day 12 (right). Scale bars, 200 μm. (C) Histological score of colon sections at day 7 and 12. One representative experiment out of two. (D) Gene Ontology analysis of microarray data showing downregulation of the expression of genes involved in host defense (GO:0006952), immune response (GO:0006955) and inflammatory response (GO:0006954) (top signature, DAVID annotation) in the colon of GF *Card9*^-/-^ versus GF WT mice at day 7 of colitis. (E) *Il-22*, *Reg3β* and *Reg3γ*, and (F) *Il-1β* and *TNF-α* expression by qRT-PCR in total colon tissue of DSS-exposed GF WT and GF *Card9*^-/-^ mice at day 0, 7 and 12, normalized to *Gapdh*. Data points represent individual mice. Data are mean ± SEM of two independent experiments. *P<0.05, **P<0.01, ***P<0.001 and ****P<0.0001, as determined by as determined by two-way analysis of variance (ANOVA) with Sidak’s post-test (A, C) and Mann-Whitney test (E, F).

### Adult *Card9*^-/-^ mice susceptibility to colitis is not overridden by colonization with a WT microbiota

To compare the impact of *Card9* deletion and the gut microbiota on colitis susceptibility, we colonized adult GF *Card9*^-/-^ or GF WT mice with the microbiota of WT mice (WT → GF *Card9*^-/-^ and WT → GF WT) and exposed them to DSS (Fig. 2A). In parallel, GF WT and GF *Card9*^-/-^ mice colonized with the microbiota of *Card9*^-/-^ mice (*Card9*^-/-^ → GF WT and *Card9*^-/-^ → GF *Card9*^-/-^ were used as controls (Fig. 2A). In accordance with our previous work, *Card9*^-/-^ → GF WT exhibited higher susceptibility to DSS-induced colitis than WT → GF WT, with impaired recovery (Fig. 2B and supp Fig 2A). The increased susceptibility to colitis observed in conventional *Card9*^-/-^ mice, as evidenced by higher weight loss and greater histopathologic alterations^8,14^, was observed in *Card9*^-/-^ mice whatever the microbiota (WT → GF *Card9*^-/-^ and *Card9*^-/-^ GF *Card9*^-/-^; Fig 2B-D, supp Fig. 2A). This result suggests that a WT microbiota is not sufficient to protect adult mice from the increased susceptibility to colitis induced by a *Card9* genetic defect. We previously showed that colon tissues and mesenteric lymph nodes (MLN) of conventional *Card9*^-/-^ mice had reduced levels of IL-22, IL-17A and IL-6 compared to WT mice as well as decreased colonic expression of *Reg3β* and *Reg3γ* after administration of DSS^8,14^. Similarly, colonic expression of *Il-22, Reg3β* and *Reg3γ* was decreased in both WT → GF *Card9*^-/-^ and *Card9*^-/-^ GF *Card9*^-/-^ mice compared to WT → GF WT (supp Fig. 2B), while no difference was observed between *Card9*^-/-^ GF *Card9*^-/-^ and WT → GF *Card9*^-/-^ mice (Fig. 2E). We also found similar amounts of IL-22, IL-17A, IL-6 and IL-10 in the colon (Fig. 2F) and MLN (Fig. 2G) of WT → GF *Card9*^-/-^ and *Card9*^-/-^ → GF *Card9*^-/-^ mice, suggesting that the genetic effect of *Card9* deletion on intestinal immune response persists in the presence of a WT microbiota. Altogether, these results demonstrate that colonization with a WT microbiota is not sufficient to rescue the increased colitis susceptibility of *Card9*^-/-^ mice, highlighting that *Card9* gene deletion intrinsically contributes to the exacerbated intestinal inflammation by altering intestinal expression and/or production of pro-inflammatory cytokines and antimicrobial peptides.

**Figure 2.**
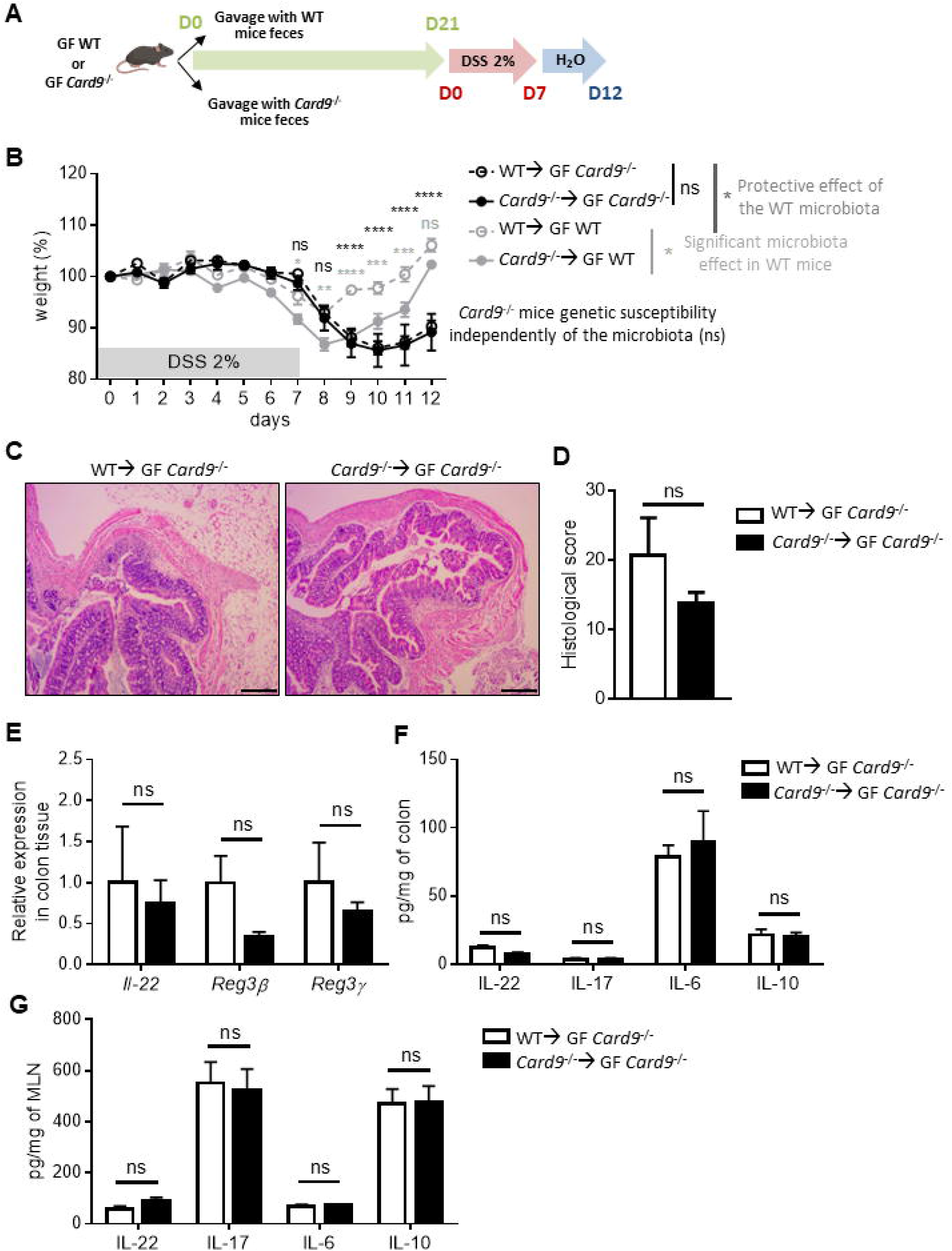
*Card9*^-/-^ mice susceptibility to colitis is not overridden by colonization with a WT microbiota. (A) Schematic representation of the DSS-induced colitis experiment preceded by 3 weeks of oral administration (gavage) of GF WT and GF *Card9*^-/-^ mice with the microbiota of WT mice (WT → GF WT and *Card9*^-/-^ → GF WT), or *Card9*^-/-^ mice (WT → GF *Card9*^-/-^ and *Card9*^-/-^ → GF *Card9*^-/-^). DSS 2% in drinking water for 7 days and then water for 5 days. (B) Weight of DSS-exposed GF *Card9*^-/-^ or GF WT mice colonized with the microbiota of WT mice (WT → GF *Card9*^-/-^ and WT → GF WT) or *Card9*^-/-^ mice (*Card9*^-/-^ → GF WT and *Card9*^-/-^ → GF *Card9*^-/-^). (C) Representative H&E-stained images of colon cross-sections from DSS-exposed GF *Card9*^-/-^ colonized with the microbiota of WT mice (WT → GF *Card9*^-/-^, left) or *Card9*^-/-^ mice (*Card9*^-/-^ → GF *Card9*^-/-^, right) at day 12. Scale bars, 500 μm. (D) Histological score of colon sections at day 12. (E) *Il-22*, *Reg3β* and *Reg3γ* expression by qRT-PCR in total colon tissue at day 12, normalized to *Gapdh*. IL-17, IL-6, IL-10 and IL-22 concentration measured by ELISA in total colon tissue (F) or mesenteric lymph nodes (MLN, G) of WT → GF *Card9*^-/-^ and *Card9*^-/-^ → GF *Card9*^-/-^ mice at day 12. Data points represent individual mice. Data are mean ± SEM. *P<0.05, **P<0.01 and ***P<0.001, as determined by two-way analysis of variance (ANOVA) with Sidak’s post-test (B) and Mann-Whitney test (D, E, F, G).

### WT microbiota shaped by *Card9* gene deletion exhibits altered composition and AhR activity

The defect in the intestinal immune response induced by *Card9* gene deletion raises the question of *Card9*^-/-^ microbiota’s contribution to the hyper-susceptibility of WT → GF *Card9*^-/-^ mice to colitis. We explored the bacterial composition of GF WT and *Card9*^-/-^ mice colonized with a WT microbiota for three weeks. Beta diversity analysis revealed major differences between the microbiota of WT → GF WT and WT → GF *Card9*^-/-^ mice throughout the experiment. The differences were already present at day 7 after microbiota colonization, but increased at day 21 (Fig. 3B). Pairwise distance and richness measurement (observed ASVs) confirmed the increased shift of microbiota composition between WT → GF WT and WT → GF *Card9*^-/-^ mice at day 21 compared to day 7 (Fig. 3A and C).Using the linear discriminant analysis effect size (LEfSe) pipeline to compare the microbiota of WT → GF WT and WT → GF *Card9*^-/-^ mice, we detected twice as many differentially represented taxa at day 21 (n=41) than at day 7 (n=22) post-colonization (Supp. Fig. 3A and B). As the colonic expression of *Il-22* and its target genes *Reg3γ* and *Reg3β* were decreased in WT → GF *Card9*^-/-^ mice compared to WT → GF WT, we reasoned that it might be related to an impaired ability of the microbiota to activate AhR and downstream IL-22 production. Using an AhR reporter system, we found that feces from WT → GF *Card9*^-/-^ mice were defective in their ability to activate AhR as rapidly as six days after colonization, similar to feces from *Card9*^-/-^ GF WT and *Card9*^-/-^ GF *Card9*^-/-^ mice (Fig. 3D and supp. Fig. 3C). These results suggest that *Card9* gene deletion contributes to the colitis susceptibility of the mice by both impairing the intestinal immune response and altering the gut microbiota composition and capacity to produce AhR ligands.

**Figure 3.**
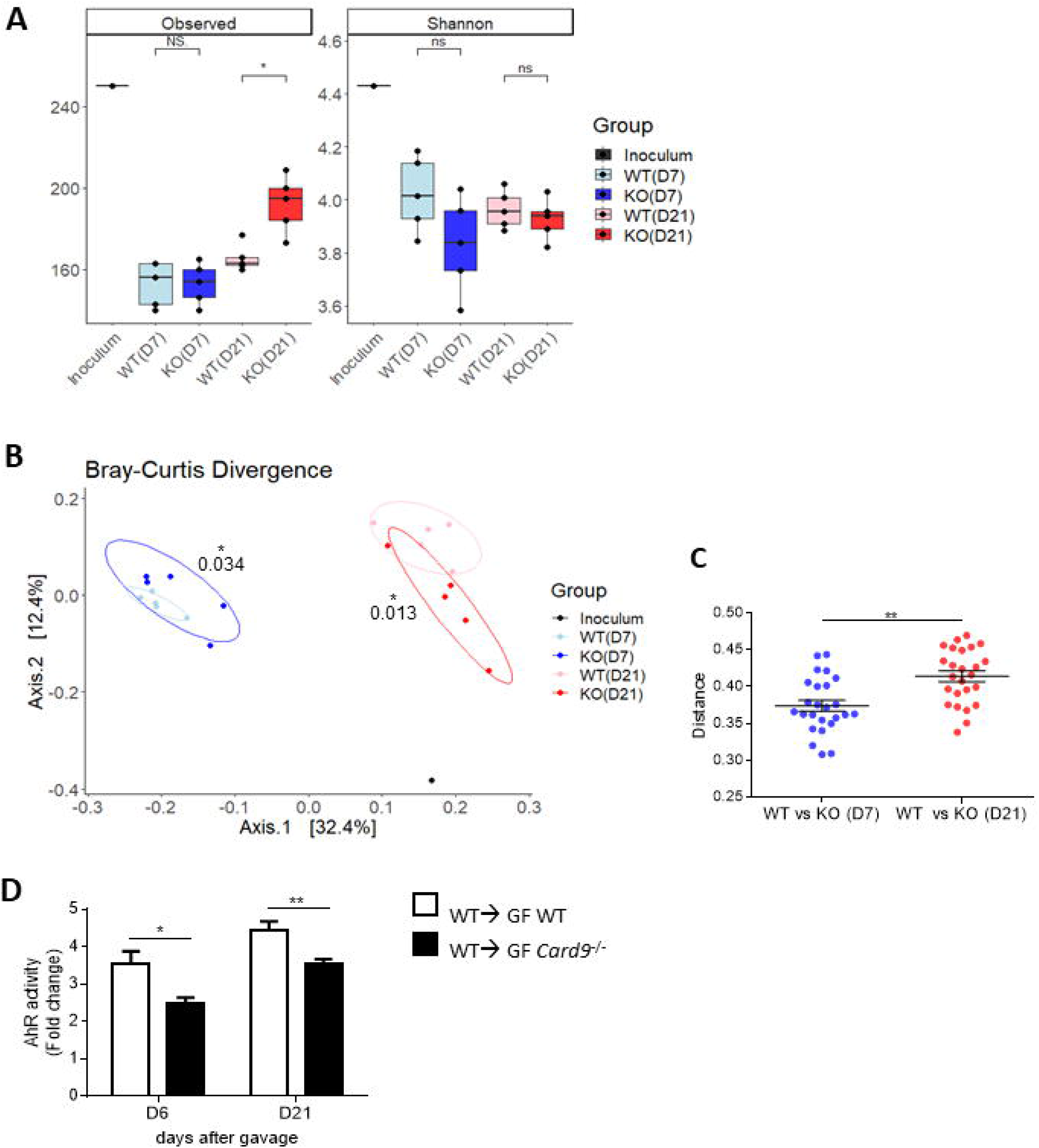
WT microbiota shaped by *Card9* gene deletion exhibits altered composition and AhR activity. (A) Alpha diversity analysis (left panel: observed ASVs, right panel: Shannon index) of the WT inoculum and the fecal microbiota of WT → GF WT and WT → GF *Card9*^-/-^ mice at day 7 and 21. (B) Beta diversity analysis (Bray-Curtis Divergence) using Jaccard index (binary) and PERMANOVA. (C) Pairwise distance (Bray-Curtis Divergence) of the fecal microbiota of WT → GF WT and WT → GF *Card9*^-/-^ mice at day 7 versus 21. (D) AhR activity (shown as ‘Fold change’) measured in feces of WT → GF WT and WT → GF *Card9*^-/-^ mice at day 6 and 21 after colonization. Data are mean ± SEM. *P<0.05 and **P<0.01, as determined by Wilcoxon test of Pairwise (C) and unpaired t test (D).

### *Card9* regulates *Lactobacillus* strains capacity to produce AhR ligands

To investigate further how *Card9* gene deletion modulates the microbiota capacity to produce AhR ligands, we colonized GF WT and GF *Card9*^-/-^ mice with the following five intestinal bacterial strains from the Proteobacteria, Bacteroidetes and Firmicutes phyla: *Escherichia coli* MG1655 and *Bacteroides thetaiotaomicron* VPI-5482 that do not produce AhR ligands, and three *Lactobacillus* strains known to produce AhR ligands, *L. murinus* CNCM I-5020, *L. reuteri* CNCM I-5022 and *L. taiwanensis* CNCM I-5019^8^ (Fig. 4A). All the microorganisms rapidly and steadily colonized the intestine of GF WT and GF *Card9*^-/-^ mice (Fig. 4B-C). However, the level of the *B. thetaiotaomicron* was lower in GF *Card9*^-/-^ compared to WT mice from the third week of colonization, suggesting some effect of the genotype on the intestinal microorganisms (Fig. 4B). Among the *Lactobacillus* strains, the level of *L. murinus* was higher than the other *Lactobacillus* strains in both genotype groups (Supp Fig. 4A). Although the *Lactobacillus* levels were not different between the groups (Fig. 4C), the capacity of the microbiota to activate AhR was decreased in GF *Card9*^-/-^ mice starting at day 11 (Fig. 4D), suggesting that CARD9 can affect the microbiota functions even without noticeable changes in the abundance of AhR agonists-producing bacteria. AhR ligands produced by the gut microbiota are known to promote IL-22 production by several intestinal immune cells, including innate lymphoid cells (ILC), T helper 17 (Th17) and Th22 cells, γδ T cells, and lymphoid tissue inducer (LTi) cells^15^. Before any microbiological contact (day 0), immune cells from the colon *lamina propria* of GF WT and GF *Card9*^-/-^ mice produced similar amounts of IL-22 and IL-17 (Fig. 4E and supp. Fig. 4B and C). Following three weeks of colonization with the five intestinal bacterial strains (day 24), we observed that IL-22 production by Th22 (CD4^+^ αβ T cells) and NKp46^+^ ILCs was decreased in the colon *lamina propria* of GF *Card9*^-/-^ mice, as compared to GF WT mice (Fig. 4E). In contrast, no difference was observed regarding IL-17 production (Supp. Fig. 4C). These data indicate that *Card9* gene deletion, without altering the population levels of *Lactobacillus*, impaired their capacity to produce AhR ligands, which induces a decreased production of IL-22 by T cells and ILCs in the colon.

**Figure 4.**
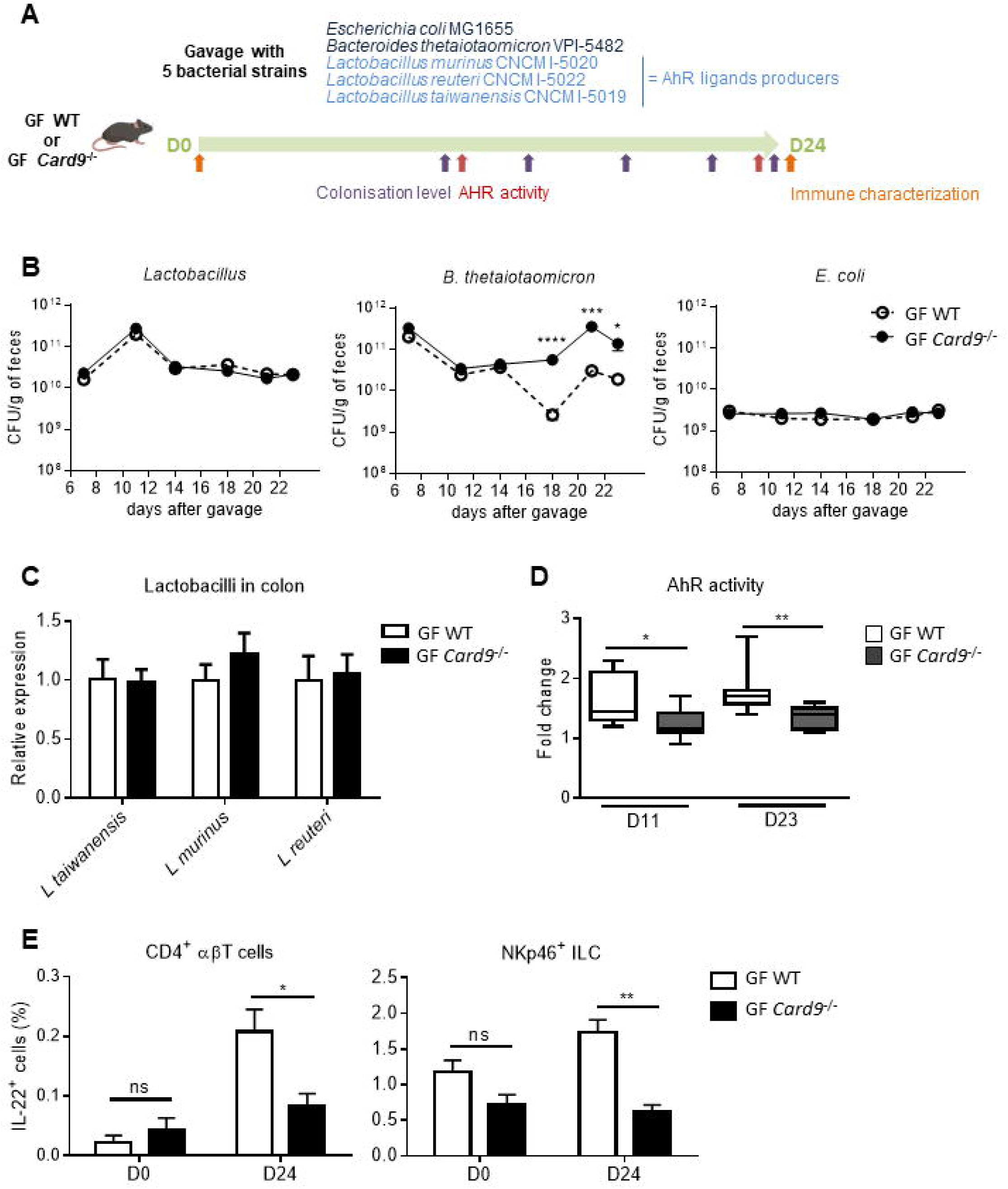
*Card9* regulates *Lactobacillus* strains capacity to produce AhR ligands. (A) Schematic representation of the gavage of GF WT or *Card9*^-/-^ mice five different bacterial strains and analyses performed. (B) Concentration of bacterial strains in feces reported as CFU/g of feces of GF WT and GF *Card9*^-/-^ mice after gavage with: three *Lactobacillus* strains known to produce AhR ligands, *L. murinus* CNCM I-5020, *L. reuteri* CNCM I-5022 and *L. taiwanensis* CNCM I-5019 (left panel); *Bacteroides thetaiotaomicron* VPI-5482 (middle panel); or *Escherichia coli* MG1655 (right panel). (C) qRT-PCR detection of each of the three gavaged *Lactobacillus* strains (*L. murinus* CNCM I-5020, *L. reuteri* CNCM I-5022 and *L. taiwanensis* CNCM I-5019) in colon of GF WT and GF *Card9*^-/-^ mice. (D) Induction of AhR activity (shown as ‘Fold change’) induced by *Lactobacillus* strains colonization in GF WT and GF *Card9*^-/-^ mice at day 11 and 23 after gavage. (E) Percentage of IL-22^+^ cells among CD4^+^ αβ T cells (left) and NKp46^+^ ILCs in the colon *lamina propria* of GF WT and GF *Card9*^-/-^ mice, at day 0 and 24 after gavage with the three *Lactobacillus* strains. Data points represent individual mice. Data are mean ± SEM. *P<0.05, **P<0.01, ***P<0.001, ****P<0.0001, as determined by Mann-Whitney test.

### Inherited microbiota controls Trp metabolism independently of host genotype

To better understand the impact of *Card9* deletion on microbiota metabolism, and particularly its ability to produce AhR ligands, we next investigated which of genetics or microbiota transmitted from the nursing mother was the most determinant for microbiota metabolism in offspring. We performed a cross-fostering experiment with conventional WT and *Card9*^-/-^ mice. Half of the litters were switched to a nursing mother of different genotype, leaving half of each original litter with their birth mother (WT mothers with half WT (m WT → WT p) and half *Card9*^-/-^ pups (m WT → *Card9*^-/-^ p); *Card9*^-/-^ mothers with half WT (m *Card9*^-/-^ → WT p) and half *Card9*^-/-^ pups (m *Card9^-/-^ → Card9^-/-^* p)) (Fig. 5A). Pups were weaned at week 4 and kept in separate cages according to their genotype and nursing mother until week 9. To evaluate the respective impact of inherited genetics and microbiota on tryptophan metabolism, we performed targeted metabolomics focusing on Trp metabolites in the feces and serum of the offspring at 9 weeks of age. Principal Component Analysis showed that metabolites composition of the feces and serum of the pups was best separated based on their nursing mother genotype rather than on their own genotype, revealing that the microbiota received before weaning from their nursing mother is more determinant than their own genotype (Fig. 5B). Indeed, the concentration of Trp and indole derivatives (SUM indoles, major AhR agonists produced by the gut microbiota) were significantly decreased in the feces and serum of pups raised by *Card9*^-/-^ mothers (m *Card9^-/-^ →* WT p and m *Card9^-/-^ → Card9^-/-^* p) compared to those raised by WT mothers (m WT → WT p and m WT → *Card9*^-/-^ p) 5 weeks after weaning, independently of the pups genotype (Fig. 5C-D and Supp Fig. 5A). In contrast, no difference in metabolite concentrations could significantly be explained by the pups’ genotype (Fig. 5C-D and Supp Fig. 5A). The two other Trp metabolism pathways, IDO and serotonin (5HT), were not more significantly impacted by the nursing mother genotype than by the pups’ ones (Supp Fig. 5B-C). These results highlight that the microbiota inherited from the mother has a higher impact on Trp metabolism than the pups’ own genotype, even 5 weeks after weaning. Accordingly, AhR activity of the caecum content was reduced in mice that inherited an altered microbiota (m *Card9^-/-^*) compared to those that received a normal one (m WT) (Fig. 5E). Altogether, these results show that *Card9* deletion affects microbiota metabolism and especially its capacity to produce AhR ligands, but this can be partially overcome by the implantation of a WT microbiota before the weaning period.

**Figure 5.**
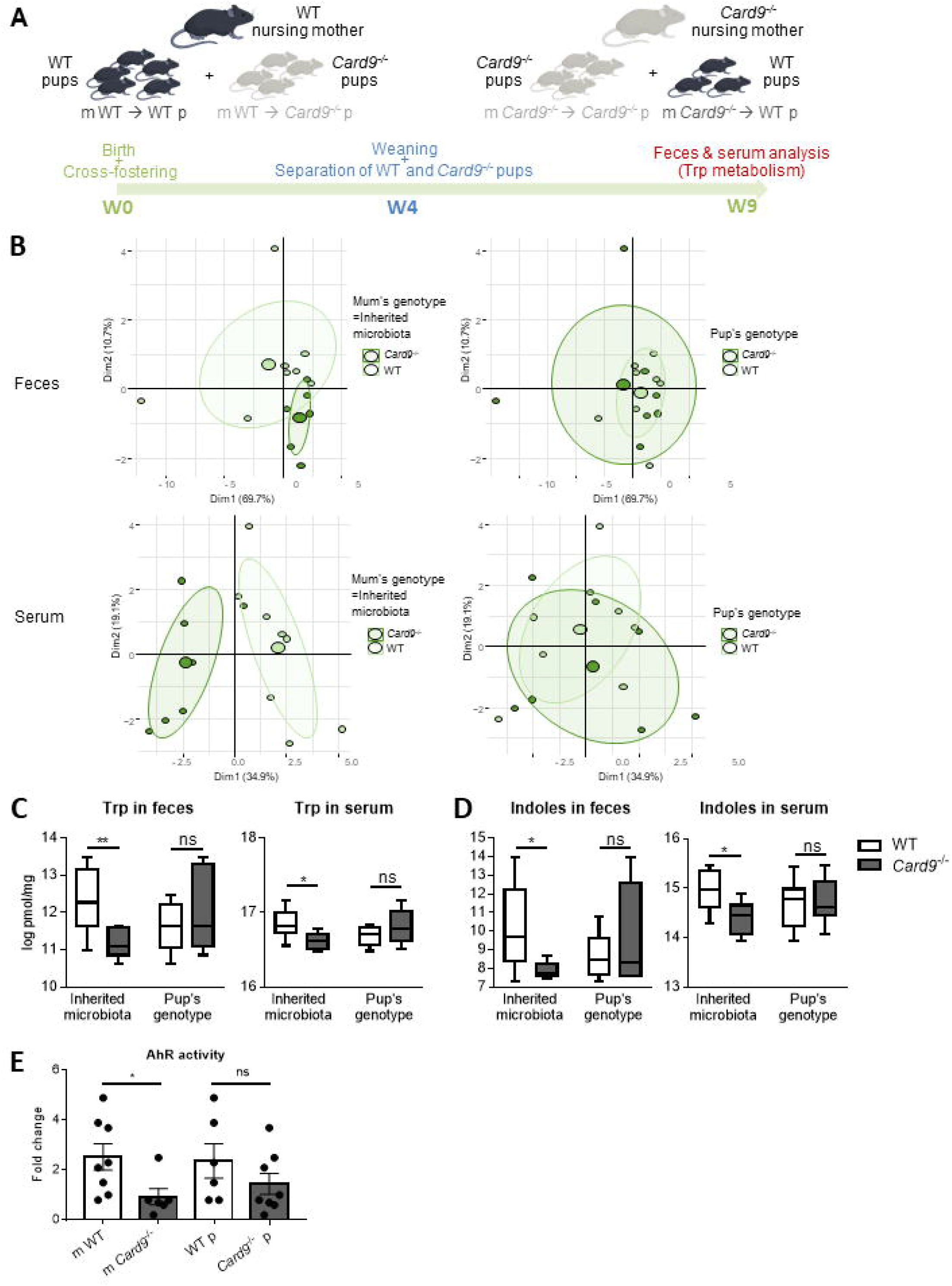
Inherited *Card9*^-/-^ microbiota controls Trp metabolism independently of the host genotype. (A) Schematic representation of the cross-fostering experiment with conventional WT and *Card9*^-/-^ mice showing the adoption of half of the offspring by a nursing mother of different phenotype, leaving half of each original litter with their birth mother (WT mothers with half WT and half *Card9*^-/-^ pups; *Card9*^-/-^ mothers with half WT and half *Card9*^-/-^ pups). Pups were weaned at week 4 and kept in separate cages according to their genotype and nursing mother until week 9, when targeted metabolomics focused on Trp metabolism on feces and serum of the pups was performed, ie 5 weeks after weaning. (B) Principal Coordinates Analysis showing metabolites composition of the feces (upper panel) and serum (lower panel) of the pups, separated according to the nursing mother genotype (left) or the pups genotype (right), at week 9 of age, i.e. 5 weeks after weaning. (C) Trp or (D) indoles concentration measured in feces (top) or serum (bottom) of the pups separated according to either the microbiota inherited from the nursing mother genotype (left) or the pups genotype (right) at week 9 of age. (E) Correlation between AhR activity induction (shown as Fold change) and functionality of *Card9* gene (in the nursing mother genotype and/or the pups genotype) at week 9 of age. Significance was determined by using spearman linear. Data points represent individual mice. *P<0.05, **P<0.01, as determined by Mann-Whitney test. Trp, tryptophan.

## Discussions

The gut microbiota is a key player in mammalian physiology, and its composition is influenced by genetics, environment, and diet ^1,2^. Any change in these factors can predispose the host to metabolic or inflammatory disorders, including obesity and IBD ^1,2^. However, it is still unclear whether dysbiosis is a cause or a consequence of these complex diseases, and it remains difficult to dissociate direct genetic effects from indirect effects through the gut microbiota. Recently, we demonstrated that the transfer of *Card9*^-/-^ mice microbiota to GF WT recipient mice was sufficient to recapitulate the defective IL-22 activation and increased colitis susceptibility observed in *Card9*^-/-^ mice ^8^. This defect, which was also observed in IBD patients with IBD-associated CARD9 SNP, is partly due to the impaired ability of the *Card9*^-/-^ microbiota to metabolize Trp into AhR ligands, such as indole derivatives ^8^. Indole derivatives, by activating AhR, regulate local IL-22 production and contribute to maintaining the fragile equilibrium between host cells and microbiota ^9–11,12^. These results directly link a functional alteration of the microbiota, i.e. altered Trp metabolism leading to reduced AhR activity and defective intestinal production of IL-22, to increased colitis susceptibility. We thus questioned the relative impacts of genotype versus microbiota on colitis susceptibility in a context of CARD9 deficiency.

We show that *Card9* modulates the susceptibility to DSS-induced colitis in both microbiota-dependent and independent manners. Using germ-free models, we proved the direct contribution of *Card9* gene in colitis recovery, through the induction of IL-22, REG3β and REG3γ, independently of the microbiota. IL-22, primarily produced by RORγt+ lymphocytes, is largely known for its critical roles in intestinal barrier function and containement of the microbiota, through the induction of antimicrobial peptides including REG3β and REG3γ^16,17^. REG3β and REG3γ contribute to the spatial segregation of intestinal bacteria and the epithelium^17–19^. Independently of microbiota-related functions, IL-22 is required for efficient mucosal wound healing via the maintenance and proliferation of epithelial stem cells^13,20^. Similarly, a growing literature points out potential metabolic functions of REG3 proteins, which could act as gut hormones^21^.

Interestingly, this direct genetic effect was not overrriden by the microbiota in our model. Indeed, a WT microbiota transplantation in adult GF *Card9*^-/-^ mice was not sufficient to compensate the genetic susceptibility, especially in terms of clinical symptoms of inflammation (weight, DAI, histology) and immune response. This is surprising as, in the other way around, the transplantation of a *Card9*^-/-^ microbiota in WT mice was sufficient to increase colitis susceptibility^8^, suggesting a dominant effect of detrimental genotype and microbiota in this model. Thus, the CARD9 protective functions against intestinal inflammation are both direct and microbiota-dependent, and rely on distinct but cumulative mechanisms.

In addition, we show that *Card9* gene deletion can shape a WT microbiota over time, with impacts on its composition and functions, especially its ability to produce AhR ligands. Indeed, we show that CARD9 modulates the AhR activity of *Lactobacillus* strains, despite the absence of impact on their colonization levels in the colon. It suggests that CARD9-dependent immune response does not reduce the abundance of *Lactobacilli in vivo*, but specifically affects their metabolic activity through a yet to determine mechanism. These data support the importance of microbiota functionality beyond taxonomic composition, in both human and mice ^22^. Indeed, these results are relevant to humans, as impaired microbial production of AhR ligands is observed in patients with IBD, and correlates with an IBD-associated SNP within CARD9 (rs10781499) ^8^. Consequently, probiotic strains that naturally produce Trp metabolites, for instance indole derivatives, could represent an interesting supportive therapy in patients with altered microbiota functions ^8,12^. However, our findings demonstrate that it is crucial to better understand the metabolic activity of these microbes *in vivo*, as they could be administered and properly colonize the colon of patients without conferring any positive effects on susceptibility to inflammation due to an altered metabolism.

Finally, a cross-fostering experiment shows that, even 5 weeks after weaning, the microbiota transmitted from the WT or *Card9*^-/-^ nursing mother (with a similar or a different genotype than the pups) has a stronger impact on the pups Trp metabolism than has the pups’ own genotype. Altogether, these results mean that *Card9* genotype modulates the microbiota metabolic capacity to produce AhR ligands, but this effect can be overridden by the implantation of a “normal” or WT microbiota before weaning. This is particularly interesting as this effect lasts up to 5 weeks after weaning, i.e. at week 9 of age, corresponding to the adulthood in mice. Previous studies showed that the reaction between the immune system and the microbiota, called “weaning reaction”, is required for immune ontology/development^23–25^. Perturbation of the weaning reaction leads to increased susceptibility to immune pathologies later in life ^23–25^. Especially, Al Nabhani and coworkers showed that the weaning reaction occurs during a specific time window, and that protection from inflammatory diseases involves microbiota-induced Treg cells ^23^. Anti-inflammatory Treg cells prevent the excessive reactivity of the immune system towards intestinal microorganisms, whereas IL-22, REG3β and REG3γ reinforce the barrier function ^17^. Moreover, it is interesting to see that, not only immune responses are impacted by the microbiota transmitted before weaning, but also feces and serum metabolism.

Owing to their tight relationship, the respective roles of host genetic factors and gut microbiota in IBD pathogenesis cannot be completely distinguished. Dysbiosis is likely both a cause and a consequence of intestinal inflammation. Our results suggest that the altered immune response in the absence of *Card9* has an effect on the microbiota composition and metabolic activity. In turn, the modified microbiota alters Trp catabolite production, affecting the host immune response and amplifying dysbiosis in a vicious cycle that leads to the loss of intestinal homeostasis. To conclude, better understanding of the impact of immune genes on microbiota metabolism is key to develop efficient therapeutic strategies for patients suffering from complex inflammatory disorders. Effort should be provided to elaborate strategies to prevent pathological imprinting early in life (breast-feeding, diet, pre and probiotics), in order to avoid the drift towards susceptibility to inflammatory disorders at a more advanced age, as microbiota perturbations in the pre-weaning period might have powerful long time effects.

## Materials and methods

### Mice

Card9-deficient mice (*Card9*^-/-^) on the C57BL/6J background were housed under specific pathogen-free conditions at the Saint-Antoine Research Center. GF C57BL/6J mice were bred in GF isolators at the CDTA (Transgenese et Archivage Animaux Modèles, CNRS, UPS44). Conventional mice were fed a standard chow diet R03 and GF mice were fed a diet without yeast R04 (SAFE). Animal experiments were performed according to the institutional guidelines approved by the local ethics committee of the French authorities, the ‘Comité d’Ethique en Experimentation Animale’ (COMETHEA) and registered with the following national number C2EA-24.

### Gut microbiota transfer and GF mice colonization

Fresh stool samples from WT or *Card9*^-/-^ mice (8-week-old males) were immediately transferred to an anaerobic chamber and diluted in LYHBHI (Brain-heart infusion) medium (BD Difco) supplemented with 1mg/ml cellobiose, 1mg/ml maltose, and 0.5mg/ml cysteine (Sigma-Aldrich). These fecal suspensions were used to inoculate mice. WT and *Card9*^-/-^ GF mice (4- to 5-week-old males) were randomly assigned to two groups and inoculated via oral gavage with 400 μl of fecal suspension (1:100) from the conventional wild-type (WT→GF) or *Card9*^-/-^ (*Card9*^-/-^→GF) mice and maintained in separate isolators. All colitis induction with DSS in WT→GF and *Card9*^-/-^→GF mice were performed 3 weeks after inoculation. To investigate how *Card9* gene deletion modulates the microbiota capacity to produce AhR ligands, WT and *Card9*^-/-^ GF mice (4- to 5-week-old males) were colonized with five intestinal bacterial strains (see text and figures). Bacterial suspensions (10^8^–10^9^ colony forming unit (CFU) in 400μl) were administered to mice by intragastric gavage. The microorganism population levels were determined weekly in the feces during implantation using culture methods (*E. coli:* MacConkey agar; *B. thetaiomicron*: M17 agar; Lactobacillus strains: MRS agar).

### Induction of DSS colitis

To induce colitis, mice were administered drinking water supplemented with 2% (wt./vol.) dextran sulfate sodium (DSS; MP Biomedicals) for 7d, and then unsupplemented water for the next 5d.

### Cytokine quantification

After filtration through a 70μm cell strainer (BD Difco) in supplemented RPMI1640 medium (10% heat-inactivated FCS, 2mM L-glutamine, 50IU/ml penicillin, 50μg/ml streptomycin; Sigma-Aldrich), 1×10^6^ MLN cells/well were stimulated by 50ng/ml PMA and 1μM ionomycin (Sigma-Aldrich) for 48h (37°C, 10%CO2). To measure cytokine levels in explants, tissues from the medial colon were rinsed in phosphate-buffered saline (PBS, Gibco). The colonic explants were cultured (37°C, 10%CO2) overnight in 24-well tissue culture plates (Corning) in 1ml of complete RPMI1640 medium. ELISAs were performed for IL-10, IL-17A (Mabtech), IL-22 (eBioscience), and IL-6 (R&D Systems). Normalization to the dry weight of each colonic explant was done.

### Lamina propria cell isolation and flow cytometry

Cells from the colon and small intestine lamina propria were isolated, stimulated and stained as previously described^8^ with specified antibodies (**supplemental table S1**). The cells were analyzed using a Gallios flow cytometer (Beckman Coulter) and identified as macrophages (MHCII+F4/80+CD103-CD11b+CD11c-), dendritic cells (MHCII+F4/80-CD103+/-CD11b-CD11c+), TH17 cells (CD3+CD4+ CD8α-IL-17+IL-22+), TH22 cells (CD3+CD4+ CD8α-IL-17-IL-22+), NKp46+ ILCs (including ILC3 and NK cells; CD3-CD4-CD8α-NKp46+), LTi cells (CD3-CD4+NKp46-), γδ T cells (CD3+CD4-CD8α-TCRγδ+) or CD3-CD4-CD8α-NKp46-cells.

### Histology

Colon samples were fixed, embedded in paraffin and stained with H&E. Slides were scanned and analysed to determine the histological score as previously described^14^.

### Gene expression analysis using quantitative reverse-transcription PCR (qRT-PCR)

Total RNA was isolated from colon samples or cell suspensions using RNeasy Mini Kit (Qiagen), and quantitative RT-PCR performed using SuperScript II Reverse Transcriptase (Life Technologies) and then a Takyon SYBR Green PCR kit (Eurogentec) in a StepOnePlus apparatus (Applied Biosystems) with specific mouse oligonucleotides (**supplemental table S1**). We used the 2-ΔΔCt quantification method with mouse Gapdh as an endogenous control and the GF WT or WF→GF WT or WT→GF *Card9*^-/-^ group as a calibrator.

### Fecal DNA extraction, bacterial quantification and AHR activity measurment

Fecal DNA was extracted from the weighted stool samples as previously described^8^. DNA was subjected to qPCR by using TaqMan Gene Expression Assays (Life Technologies) for quantification of all bacterial sequences (see probes and primers for the bacterial 16S rDNA genes in **supplemental table 1**). The CT–ΔΔCt method was used. The expression level of eahc individual Lactobacilli species was expressed relative to “All Lactobacilli”. Luciferase assay for AHR activity measurement was performed as previously described^8^.

### 16S rDNA gene sequencing and analysis

DNA was isolated from the feces of mice as previously described^8^.

Paired-end sequences were analysed using the *dada2^26^* (v1.24.0) algorithm in the R statistical programming environment (v4.2.0, 2018) to produce amplicon sequence variants (ASVs). Taxonomic assignment was performed using the SILVA reference database^27^ (v138.1). Alpha diversity were presented using the Shannon index and number of observed ASVs. Beta diversity was calculated using the binary Jaccard distance and plotted using principle coordinate analyses (PCoA) plots in the *vegan* package (v2.6-2). Groupings were tested using PERMANOVA with 999 permutations and the ‘adonis’ function for each timepoint separately. Differential abundance testing was performed using the linear discriminant analysis with effect size (LEfSe) on proportional (total sum normalised) data, using default settings^28^. Plotting was performed in *ggplot2* (v3.3.6) and *ggpubr* (v0.4.0), using the Wilcoxon rank sum test. Analysis scripts are available on github (https://github.com/ajlavelle/dyscolic-Figures). Sequences are deposited on sequence read archive (accession number pending).

### Gene expression by microarray analyses

Total RNA was isolated using the protocol described above. RNA integrity was verified using a Bioanalyser2100 with RNA6000 Nano chips (Agilent Technologies). Transcriptional profiling was performed on mouse colon samples using the SurePrint G3 Mouse GE8×60K Microarray kit (design ID:028005, Agilent Technologies). Cyanine-3 (Cy3)-labeled cRNAs were prepared with 100ng of total RNA using a One-Color Low Input Quick Amp Labeling kit (Agilent Technologies). The specific activities and cRNA yields were determined by using a NanoDrop ND-1000 (Thermo Fisher Scientific). For each sample, 600 ng of Cy3-labeled cRNA (specific activity> 11.0 pmol Cy3/μg of cRNA) were fragmented at 60°C for 30min and hybridized to the microarrays for 17h at 65°C in a rotating hybridization oven (Agilent Technologies). After hybridization, the microarrays were washed and then immediately dried. After washing, the slides were scanned using a G2565CA Scanner System (Agilent Technologies) at a resolution of 3μm and a dynamic range of 20bits. The resulting TIFF images were analyzed using the Feature Extraction Software v10.7.3.1 (Agilent Technologies) according to the GE1_107_Sep09 protocol. The microarray data were submitted to GEO under accession number (pending).

### Microarray analysis

The R package ‘agilp’ was used to pre-process the raw data. Agilent Feature Extraction software computed a P value for each probe in each array to test whether the scanned signals were significantly higher than the background signal. Probes were considered to be detected if the P value was <0.05, and if at least 60% of samples per group and under at least one condition. After normalization using Quantile normalization, spike-in, positive and negative control probes were removed from the normalized data. For differential expression analysis, we used the limma eBayes test. The Benjamini– Hochberg correction method was used to control the false-discovery rate (FDR). All significant gene lists were annotated for enriched biological functions and pathways using the DAVID platform^29^ for gene ontology (GO) and Kyoto Encyclopedia of Genes and Genomes (KEGG) terms. Significant canonical pathways had adjusted P values, according to Benjamini’s method, below 0.05. DAVID was performed to test for the biological pathway enrichment of Venn’s elements.

### Cross-fostering experiment

Breeding pairs of conventional WT and *Card9*^-/-^ mice were set up simultaneously to obtain synchronized births. Half of the litters were switched to a nursing mother of different genotype, leaving half of each original litter with their birth mother (WT mothers with half WT and half *Card9*^-/-^ pups; *Card9*^-/-^ mothers with half WT and half *Card9*^-/-^ pups) (see **Fig. 5A**). Pups were weaned at week 4 and kept in separate cages according to their genotype and nursing mother until week 9.

### Targeted metabolomics

Samples were lyophilized and weighted. The methods has been described previously^30^. Briefly, internal standard (100μL) and methanol/water (50:50) were added in each samples. Supernatant were collected after an agitation during 30min at 4°C and centrifugation. After simultaneous evaporation, each well was resuspended in 100μL of a methanol/water mixture (1:9). Finally, 5μL were injected into the LC-MS (XEVO-TQ-XS, Waters®). A Kinetex C18 xb column (1.7μm × 150mm × 2.1mm, temperature 55°C) associated with a gradient of two mobile phases (Phase A:Water + 0.5% formic acid; Phase B: MeOH + 0.5% formic acid) at a florate of 0.4mL/min was used. A calibration curve was created by calculating the intensity ratio obtained between each metabolite and its internal standard. These calibration curves were then used to determine the concentrations of each metabolite in patient samples.

### Statistical analyses

GraphPad Prism version 6.0 was used for all analyses using statistical tests specified in figure legends.

## Declaration section

### Ethics approval and Consent to participate

All authors consent to participate in this study

### Consent for publication

All authors give their consent for publication

### Availability of data and materials

All data related to this study will be made available to other researchers: GEO for microarray, SRA for 16s RNA sequences and analysis (accession numbers pending).

### Authors’ contributions

C. D., B.L. and H.S. conceived and designed the study, performed data analysis, and wrote the manuscript; C.D. and B.L. designed and conducted all experiments, unless otherwise indicated; A.La. performed the analysis of the microbiota composition; A.Le. and P.E. realized the targeted metabolomics analyses; H.-P.P. and M.B. conducted the bioinformatics studies and analyzed the microarray experiments; M.S. and performed the AhR activity experiments; M.-L.M., G.D.C., C.B. and J.P. provided technical help for the in vitro and in vivo experiments; C.D., B.L., P.L. and H.S. discussed the experiments and results.

## Acknowledgements

We thank the members of the IERP (INRAE) and PHEA (CRSA) animal facilities, and of the @bridge histology platform (Université Paris-Saclay, INRAE, AgroParisTech, GABI) for their contribution to this study.

## Supplemental information

**Supp Figure 1.** Genes included in the Gene Ontology pathways downregulated in the colon of GF *Card9*^-/-^ versus GF WT mice at day 7 of colitis (host defense (GO:0006952), immune response (GO:0006955) and inflammatory response (GO:0006954)), including *Reg3β* and *Il1rl1*.

**Supp Figure 2.** (A) Disease activity index (DAI) of DSS-exposed GF *Card9*^-/-^ or GF WT mice colonized with the microbiota of WT or *Card9*^-/-^ mice. For statistical comparisons, † indicates WT → GF WT versus WT → GF *Card9*^-/-^, ‡ indicates *Card9*^-/-^ → GF WT versus *Card9*^-/-^ → GF *Card9*^-/-^ and * indicates WT → GF WT versus *Card9*^-/-^ → GF WT. (B) *Il-22*, *Reg3β* and *Reg3γ* expression by qRT-PCR in total colon tissue of WT → GF WT, WT → GF *Card9*^-/-^ and *Card9*^-/-^ → GF *Card9*^-/-^ mice at day 12, normalized to *Gapdh*. Data are mean ± SEM. *P<0.05, †P<0.05, ‡ P<0.05, **P<0.01, ‡‡P<0.01, ***P<0.001 and †††P<0.001 as determined by one way ANOVA and post hoc Tukey test (A) and Mann-Whitney test (B).

**Supp Figure 3.** LEFse analyses showing taxa (genus level) overrepresented (positive values, right) and underrepresented (negative values, left) in the microbiota of WT → GF WT compared to WT → GF *Card9*^-/-^ mice at day 7 (A) and 21 (B). (C) AHR activity (shown as fold change) of feces of WT → GF WT, *Card9*^-/-^ → GF WT, WT → GF *Card9*^-/-^ and *Card9*^-/-^ → GF *Card9*^-/-^ mice at day 6 and 21. Data are mean ± SEM. *P<0.05 and **P<0.01, as determined by Mann-Whitney test.

**Supp. Figure 4.** (A) Relative expression of each of the three gavaged *Lactobacillus* strains (*L. murinus* CNCM I-5020, *L. reuteri* CNCM I-5022 and *L. taiwanensis* CNCM I-5019) in feces of GF WT (left) and GF *Card9*^-/-^ mice (right), assessed by qRT-PCR and normalized to “All lactobacilli” quantity at day 24. (B) Percentage of IL-22^+^ cells among all immune cells, γδT cells DN, CD4^+^ ILCs, and NKp46^-^ CD4^-^ ILCs in the colon *lamina propria* of GF WT and GF *Card9*^-/-^ mice, at day 0 and 24 after gavage with the three *Lactobacillus* strains. (C) Percentage of IL-17^+^ cells among all cells, γδT cells DN, CD4^+^ ILCs, NKp46^-^ CD4^-^ ILCs, CD4^+^ αβT cells and NKp46^+^ ILCs in the colon *lamina propria* of GF WT and GF *Card9*^-/-^ mice, at day 0 and 24 after gavage with the three *Lactobacillus* strains. Data points represent individual mice. Data are mean ± SEM. *P<0.05, **P<0.01, ***P<0.001, as determined by Mann-Whitney test. DN, double negative.

**Supp Figure 5.** (A) Indole 3 lactic acid concentration in feces of the pups separated according to the nursing mother genotype or the pups’ genotype 5 weeks after weaning. Metabolites concentration from the IDO (B) and serotonin (5HT, C) pathways measured in feces or serum of the pups separated according to the nursing mother genotype or the pups’ genotype, 5 weeks after weaning. Data points represent individual mice. *P<0.05, as determined by Mann-Whitney test. Trp, tryptophan.

**Supplemental table 1**. Antibodies and nucleotides list.

## References

1. Ananthakrishnan, A.N. (2015). Epidemiology and risk factors for IBD. Nat Rev Gastroenterol Hepatol 12, 205–217. 10.1038/nrgastro.2015.34.

2. Silva, M.J.B., Carneiro, M.B.H., dos Anjos Pultz, B., Pereira Silva, D., Lopes, M.E. de M., and dos Santos, L.M. (2015). The Multifaceted Role of Commensal Microbiota in Homeostasis and Gastrointestinal Diseases. Journal of Immunology Research 2015, 1–14. 10.1155/2015/321241.

3. Levy, M., Kolodziejczyk, A.A., Thaiss, C.A., and Elinav, E. (2017). Dysbiosis and the immune system. Nat Rev Immunol 17, 219–232. 10.1038/nri.2017.7.

4. Hsu, Y.-M.S., Zhang, Y., You, Y., Wang, D., Li, H., Duramad, O., Qin, X.-F., Dong, C., and Lin, X. (2007). The adaptor protein CARD9 is required for innate immune responses to intracellular pathogens. Nat Immunol 8, 198–205. 10.1038/ni1426.

5. Lanternier, F., Mahdaviani, S.A., Barbati, E., Chaussade, H., Koumar, Y., Levy, R., Denis, B., Brunel, A.-S., Martin, S., Loop, M., et al. (2015). Inherited CARD9 deficiency in otherwise healthy children and adults with Candida species–induced meningoencephalitis, colitis, or both. Journal of Allergy and Clinical Immunology 135, 1558–1568.e2. 10.1016/j.jaci.2014.12.1930.

6. Goodridge, H.S., Shimada, T., Wolf, A.J., Hsu, Y.-M.S., Becker, C.A., Lin, X., and Underhill, D.M. (2009). Differential use of CARD9 by dectin-1 in macrophages and dendritic cells. J Immunol 182, 1146–1154. 10.4049/jimmunol.182.2.1146.

7. Hara, H., Ishihara, C., Takeuchi, A., Imanishi, T., Xue, L., Morris, S.W., Inui, M., Takai, T., Shibuya, A., Saijo, S., et al. (2007). The adaptor protein CARD9 is essential for the activation of myeloid cells through ITAM-associated and Toll-like receptors. Nat Immunol 8, 619–629. 10.1038/ni1466.

8. Lamas, B., Richard, M.L., Leducq, V., Pham, H.-P., Michel, M.-L., Da Costa, G., Bridonneau, C., Jegou, S., Hoffmann, T.W., Natividad, J.M., et al. (2016). CARD9 impacts colitis by altering gut microbiota metabolism of tryptophan into aryl hydrocarbon receptor ligands. Nat Med 22, 598–605. 10.1038/nm.4102.

9. Rutz, S., Eidenschenk, C., and Ouyang, W. (2013). IL-22, not simply a Th17 cytokine. Immunol Rev 252, 116–132. 10.1111/imr.12027.

10. Sonnenberg, G.F., Fouser, L.A., and Artis, D. (2011). Border patrol: regulation of immunity, inflammation and tissue homeostasis at barrier surfaces by IL-22. Nat Immunol 12, 383–390. 10.1038/ni.2025.

11. Lee, J.S., Cella, M., McDonald, K.G., Garlanda, C., Kennedy, G.D., Nukaya, M., Mantovani, A., Kopan, R., Bradfield, C.A., Newberry, R.D., et al. (2012). AHR drives the development of gut ILC22 cells and postnatal lymphoid tissues via pathways dependent on and independent of Notch. Nat Immunol 13, 144–151. 10.1038/ni.2187.

12. Zelante, T., Iannitti, R.G., Cunha, C., De Luca, A., Giovannini, G., Pieraccini, G., Zecchi, R., D’Angelo, C., Massi-Benedetti, C., Fallarino, F., et al. (2013). Tryptophan Catabolites from Microbiota Engage Aryl Hydrocarbon Receptor and Balance Mucosal Reactivity via Interleukin-22. Immunity 39, 372–385. 10.1016/j.immuni.2013.08.003.

13. Pickert, G., Neufert, C., Leppkes, M., Zheng, Y., Wittkopf, N., Warntjen, M., Lehr, H.-A., Hirth, S., Weigmann, B., Wirtz, S., et al. (2009). STAT3 links IL-22 signaling in intestinal epithelial cells to mucosal wound healing. Journal of Experimental Medicine 206, 1465–1472. 10.1084/jem.20082683.

14. Sokol, H., Conway, K.L., Zhang, M., Choi, M., Morin, B., Cao, Z., Villablanca, E.J., Li, C., Wijmenga, C., Yun, S.H., et al. (2013). Card9 Mediates Intestinal Epithelial Cell Restitution, T-Helper 17 Responses, and Control of Bacterial Infection in Mice. Gastroenterology 145, 591–601.e3. 10.1053/j.gastro.2013.05.047.

15. Lindahl, H., and Olsson, T. (2021). Interleukin-22 Influences the Th1/Th17 Axis. Front. Immunol. 12, 618110. 10.3389/fimmu.2021.618110.

16. Lo, B.C., Shin, S.B., Canals Hernaez, D., Refaeli, I., Yu, H.B., Goebeler, V., Cait, A., Mohn, W.W., Vallance, B.A., and McNagny, K.M. (2019). IL-22 Preserves Gut Epithelial Integrity and Promotes Disease Remission during Chronic Salmonella Infection. J.I. 202, 956–965. 10.4049/jimmunol.1801308.

17. Caruso, R., Lo, B.C., and Núñez, G. (2020). Host–microbiota interactions in inflammatory bowel disease. Nat Rev Immunol 20, 411–426. 10.1038/s41577-019-0268-7.

18. Vaishnava, S., Yamamoto, M., Severson, K.M., Ruhn, K.A., Yu, X., Koren, O., Ley, R., Wakeland, E.K., and Hooper, L.V. (2011). The antibacterial lectin RegIIIgamma promotes the spatial segregation of microbiota and host in the intestine. Science 334, 255–258. 10.1126/science.1209791.

19. Sonnenberg, G.F., Monticelli, L.A., Alenghat, T., Fung, T.C., Hutnick, N.A., Kunisawa, J., Shibata, N., Grunberg, S., Sinha, R., Zahm, A.M., et al. (2012). Innate lymphoid cells promote anatomical containment of lymphoid-resident commensal bacteria. Science 336, 1321–1325. 10.1126/science.1222551.

20. Lindemans, C.A., Calafiore, M., Mertelsmann, A.M., O’Connor, M.H., Dudakov, J.A., Jenq, R.R., Velardi, E., Young, L.F., Smith, O.M., Lawrence, G., et al. (2015). Interleukin-22 promotes intestinal-stem-cell-mediated epithelial regeneration. Nature 528, 560–564. 10.1038/nature16460.

21. Shin, J.H., and Seeley, R.J. (2019). Reg3 Proteins as Gut Hormones? Endocrinology 160, 1506–1514. 10.1210/en.2019-00073.

22. Heintz-Buschart, A., and Wilmes, P. (2018). Human Gut Microbiome: Function Matters. Trends in Microbiology 26, 563–574. 10.1016/j.tim.2017.11.002.

23. Al Nabhani, Z., Dulauroy, S., Marques, R., Cousu, C., Al Bounny, S., Déjardin, F., Sparwasser, T., Bérard, M., Cerf-Bensussan, N., and Eberl, G. (2019). A Weaning Reaction to Microbiota Is Required for Resistance to Immunopathologies in the Adult. Immunity 50, 1276–1288.e5. 10.1016/j.immuni.2019.02.014.

24. Gensollen, T., Iyer, S.S., Kasper, D.L., and Blumberg, R.S. (2016). How colonization by microbiota in early life shapes the immune system. Science 352, 539–544. 10.1126/science.aad9378.

25. Gollwitzer, E.S., and Marsland, B.J. (2015). Impact of Early-Life Exposures on Immune Maturation and Susceptibility to Disease. Trends in Immunology 36, 684–696. 10.1016/j.it.2015.09.009.

26. Callahan, B.J., McMurdie, P.J., Rosen, M.J., Han, A.W., Johnson, A.J.A., and Holmes, S.P. (2016). DADA2: High-resolution sample inference from Illumina amplicon data. Nat Methods 13, 581–583. 10.1038/nmeth.3869.

27. Quast, C., Pruesse, E., Yilmaz, P., Gerken, J., Schweer, T., Yarza, P., Peplies, J., and Glöckner, F.O. (2012). The SILVA ribosomal RNA gene database project: improved data processing and web-based tools. Nucleic Acids Research 41, D590–D596. 10.1093/nar/gks1219.

28. Segata, N., Izard, J., Waldron, L., Gevers, D., Miropolsky, L., Garrett, W.S., and Huttenhower, C. (2011). Metagenomic biomarker discovery and explanation. Genome Biol 12, R60. 10.1186/gb-2011-12-6-r60.

29. Sherman, B.T., Hao, M., Qiu, J., Jiao, X., Baseler, M.W., Lane, H.C., Imamichi, T., and Chang, W. (2022). DAVID: a web server for functional enrichment analysis and functional annotation of gene lists (2021 update). Nucleic Acids Research 50, W216–W221. 10.1093/nar/gkac194.

30. Lefèvre, A., Mavel, S., Nadal-Desbarats, L., Galineau, L., Attucci, S., Dufour, D., Sokol, H., and Emond, P. (2019). Validation of a global quantitative analysis methodology of tryptophan metabolites in mice using LC-MS. Talanta 195, 593–598. 10.1016/j.talanta.2018.11.094.

