## Supplementary figures and images for "Dissecting the respective roles of microbiota and host genetics in the susceptibility of *Card9*^-/-^ mice to colitis"

### Supp Figure 1

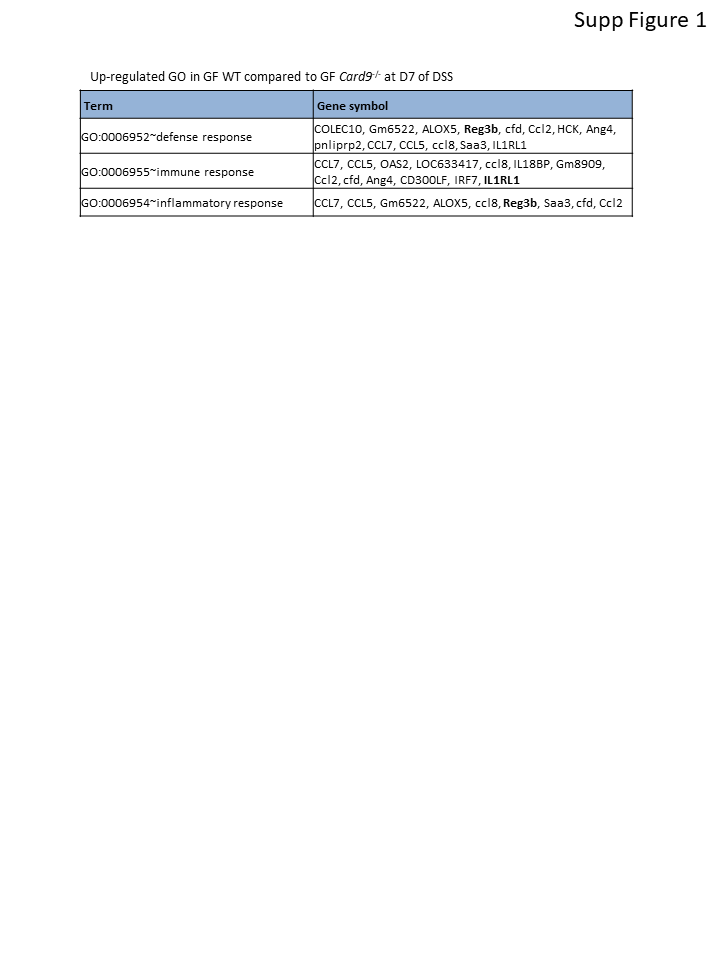

### Supp Figure 2

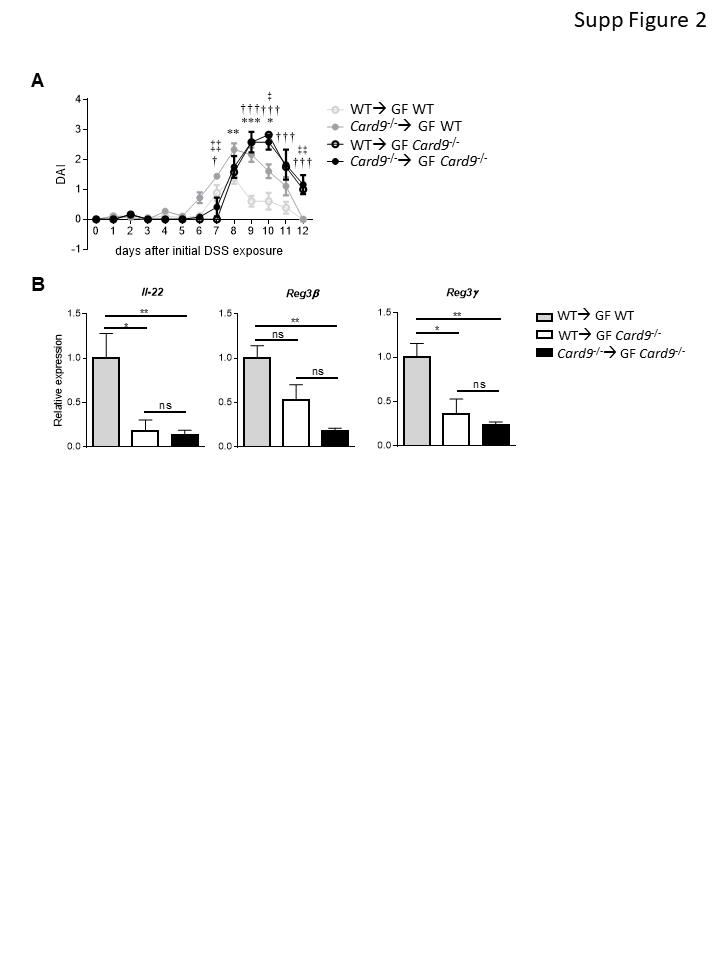

### Supp Figure 3

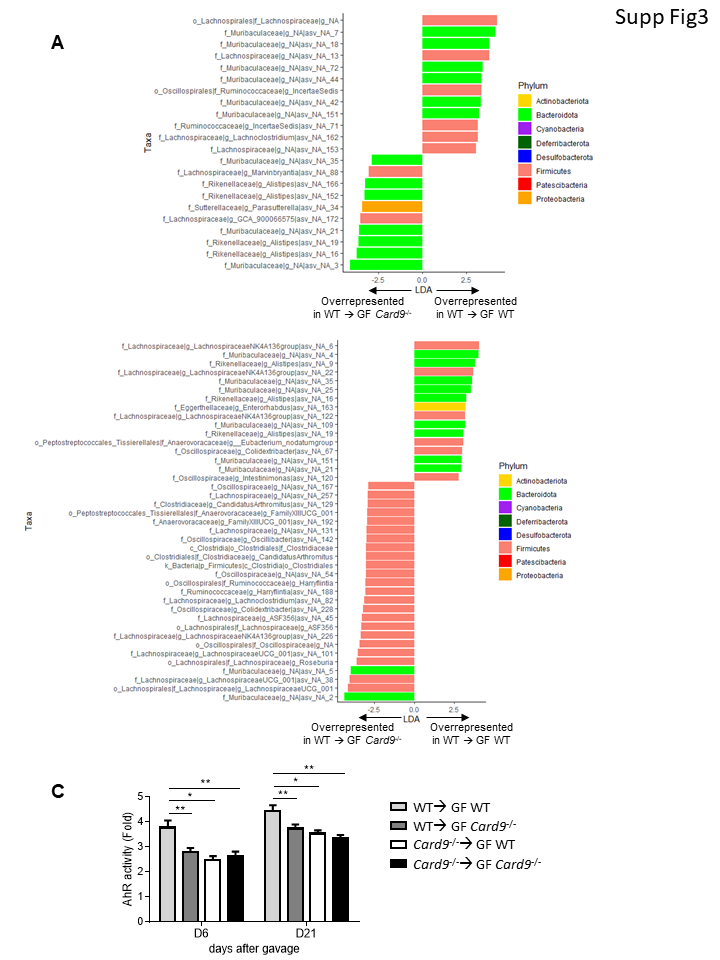

### Supp Figure 4

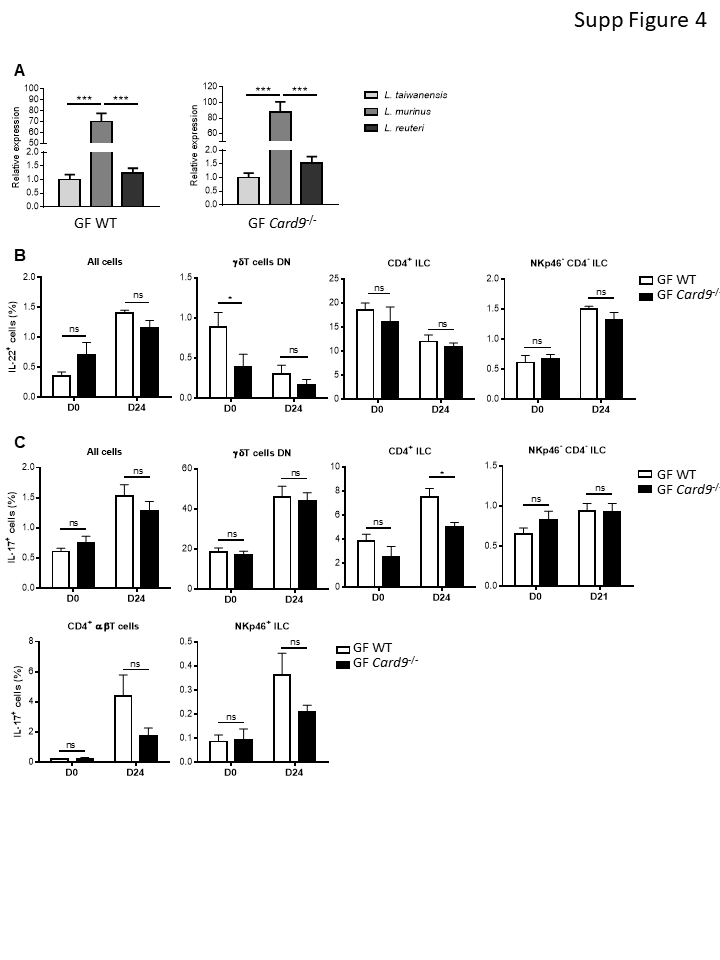

### Supp Figure 5

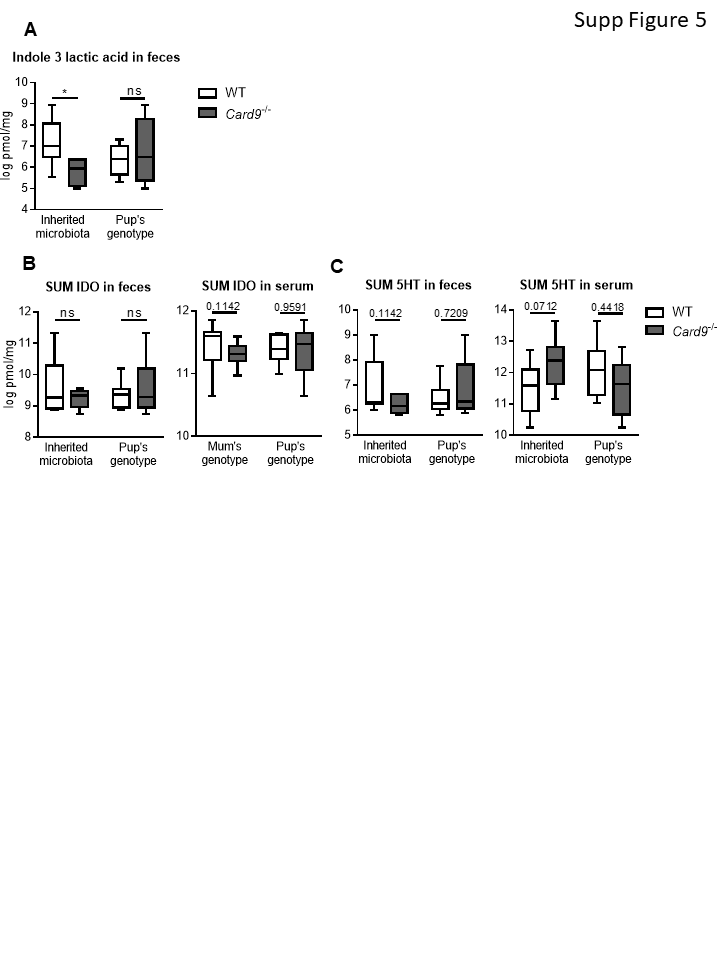
